# COX-1 – lipid interactions: arachidonic acid, cholesterol, and phospholipid binding to the membrane binding domain of COX-1

**DOI:** 10.1101/2020.05.21.109363

**Authors:** Besian I. Sejdiu, D. Peter Tieleman

## Abstract

Cyclooxygenases carry out the committed step in prostaglandin synthesis and are the target of NSAIDs, the most widely used class of drugs in alleviating pain, fever, and inflammation. While extensively studied, one aspect of their biology that has been neglected is their interaction with membrane lipids. Such lipid-protein interactions have been shown to be a driving force behind membrane protein function and activity. Cyclooxygenases (COX-1 and COX-2) are bound on the luminal side of the endoplasmic reticulum membrane. The entrance to their active site is formed by a long hydrophobic channel which is used by the cyclooxygenase natural substrate, arachidonic acid, to access the enzyme. Using atomistic and coarse-grained simulations, we show that several membrane lipids are capable of accessing the same hydrophobic channel. We observe the preferential binding of arachidonic acid, cholesterol and glycerophospholipids with residues lining the cavity of the channel. We find that the membrane binding domain (MBD) of COX-1 is usually in a lipid-bound state and not empty. This orthosteric binding by other lipids suggests a potential regulatory role of membrane lipids with the possibility of affecting the COX-1 turnover rate. We also observed the unbiased binding of arachidonic acid to the MBD of COX-1 allowing us to clearly delineate its binding pathway. We identified a series of arginine residues as being responsible for guiding arachidonic acid towards the binding site. Finally, we were also able to identify the mechanism by which COX-1 induces a positive curvature on the membrane environment.

## Introduction

Prostaglandin endoperoxide H synthase-1 and 2 isoenzymes, commonly known as cyclooxygenase-1 and 2 (COX-1 and COX-2), respectively, are membrane-bound enzymes that carry out the committed step in prostaglandin synthesis, and are thus either directly or indirectly involved in many malfunctions and pathologies, with large therapeutic implications(1–5). The anti-inflammatory, analgesic, antipyretic and antithrombotic effects of nonsteroidal anti-inflammatory drugs (NSAIDs) are associated with the inhibition of the activity of cyclooxygenases (COX-2 for the former three, and COX-1 isoform for the latter one)(6–8).

Prostanoids, the end products of cyclooxygenase (and other enzymes) catalysis, are biologically active compounds formed from arachidonic acid as the precursor. Cyclooxygenases, in a two-step process, convert arachidonate to prostaglandin H_2_ (PGH_2_): first, through the addition of two O_2_ molecules arachidonate is converted to prostaglandin G_2_ (this is the cyclooxygenase – or COX – reaction) which is then followed by a 2*e*^-^ reduction reaction to produce PGH_2_ (the peroxidase – or POX – reaction). PGH_2_ then through the action of different enzymes is converted into other prostaglandins, prostacyclin or thromboxane A_2_ (9). Structurally, COX-1 and 2 are homodimers (Figure 1), with each monomer composed of a large globular (or catalytic) domain, an epidermal growth factor-like (EGF) domain, and a membrane binding domain (MBD) (10).

**Figure 1.**
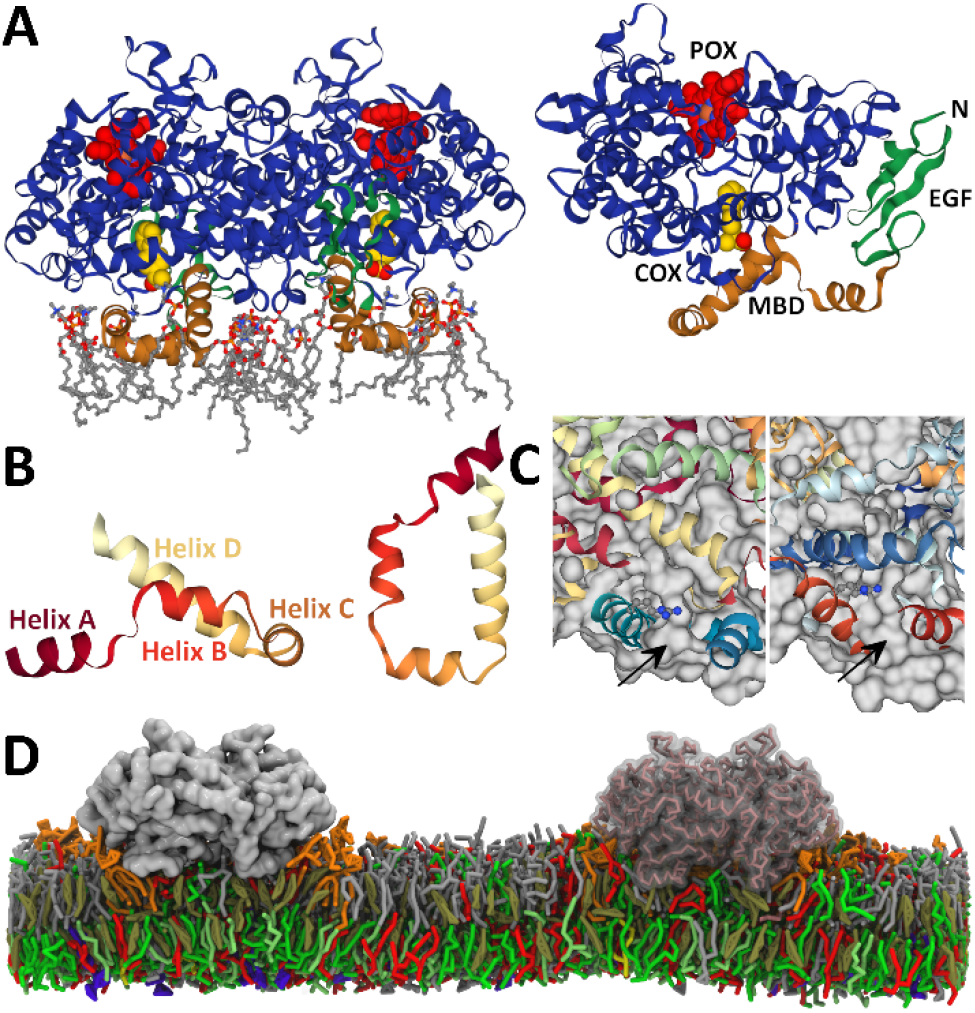
*Structure of the ovine* Prostaglandin endoperoxide H synthase-1. **A.** The enzyme is shown with its domains colored in blue, green and gold denoting the catalytic, epidermal growth factor-like and membrane binding domains, respectively. Heme is shown in red, flurbiprofen in yellow, and POPC lipids in contact with it during the simulation as stick representation. This figure is based on a similar figure from reference (2). **B.** Close-up view of the MBD of COX-1, showing the location of helices A-D, and surface rendering highlighting the hydrophobic channel formed by these helices (**C**). For reference, Arg-120 is drawn using a stick presentation. **D.** Schematic view of one of the simulated systems for COX-1, showing different lipids species using different colors. One of the proteins is drawn transparent to reveal the backbone beads as they are modeled in the MARTINI model.

COX-1 and 2 are integral monotopic membrane proteins, permanently bound to one side of the membrane without spanning the full bilayer. The MBD of COX-1 and 2 forms the entrance to a 25Å long hydrophobic channel(2) that starts at the surface of the membrane and extends to the core of the catalytic domain where it ends at the cyclooxygenase reaction site (Figure 1C). Arachidonic acid enters the enzyme via the membrane through this channel, as do NSAID drugs(11, 12). The MBD is composed of four short helical segments (helices A-D) that anchor COX-1 into the upper (outer or luminal) leaflet(13) of the endoplasmic reticulum (ER) and inner membrane of the nuclear envelope(14). While cyclooxygenases are sequence homodimers, they function as heterodimers, with one subunit serving an allosteric (E_allo_) and the other a catalytic (E_cat_) role(15, 16). This is referred to as half of site COX activity whereby only one subunit is functionally active at a given time and is regulated allosterically by the other subunit. The cross-talk between the monomers is heavily influenced by ligand binding and often results in different outcomes for COX-1 and COX-2(9).

Many aspects of cyclooxygenases have been studied, including substrate binding and enzymatic activity, kinetic profile, as well as ligand binding and inhibition(9, 11, 17). Here we focus on the lipid-protein interactions that may modulate the activity of COX-1. The study of membrane protein interactions with their lipid environment has gained a lot of popularity in recent years and has been a major focus of current research literature. Molecular Dynamics (MD) simulations, in particular, have provided many insights into the nature of these interactions, and the current understanding of lipid-protein interactions highlights the importance of direct and specific interactions with lipids, as well as general changes to the membrane physical properties in affecting the function and activity of embedded proteins(18–20). Several previous simulation studies have focused on cyclooxygenase catalytic activity(6, 7, 21, 22) and more general membrane binding of monotopic proteins(23–27). For example, the ability of dimyristoylphosphatidylcholine (DMPC) headgroup to interact with COX-1 has been noted as early as 2000(23).

To study the lipid-protein interactions COX-1 and obtain a comprehensive understanding of their interaction profile, we performed long-scale molecular dynamics (MD) simulations of COX-1 using both atomistic (AA) and coarse-grained (CG) resolutions in both simple and complex membrane environments.

The combined results from these simulations highlight the ability of lipids, such as arachidonic acid, cholesterol and glycerophospholipids, to enter into the hydrophobic channel of COX-1 and interact with hydrophobic residues lining the interior of the channel and via charge-charge interactions with Arg-120. We find two mechanisms for arachidonic acid binding, revealing thus the pathway the lipid takes to enter the active site of the enzyme. We identify several residues that are involved in guiding arachidonic acid and in maintaining its binding to the hydrophobic channel. Lastly, we also identify the mechanism by which COX-1 induces previously reported perturbations into the surrounding lipid environment that result in the creation of a strong positive curvature.

## Methods

### Protein Structure

We retrieved the protein structure from the Protein Data Bank with PDB entry 1Q4G(28). For CG simulations, all ligands – including heme – where removed before the structure was converted into a coarse-grained representation using the *martinize* tool as described on the MARTINI(29) website *(cgmartini.nl).* For atomistic simulations, heme was included in the simulations, but all glycans alongside other ligands were excluded. Table S1 provides a complete list of setups and simulation details.

### Coarse-Grained Molecular Dynamics Simulations

In terms of membrane composition, three types of systems were employed in our CG simulations: POPC only, ER-like membranes, and a membrane with complex lipid composition.

POPC membranes were used to highlight the curvature inducing ability of COX-1, whereas CG simulations with lipid concentrations mimicking closely the lipid composition of ER membranes(30–32) were used to study lipid – COX-1 interactions in multicomponent membranes. These systems contained one copy of the enzyme inserted into the membrane. For the complex membrane setup, four copies of the coarse-grained protein were placed in a 40×40 nm^2^ area which was then filled with lipids using the tool *insane(33).* It is composed of 63 different lipid types based on the model developed by Ingólfsson et al.(34) and applied to 10 different proteins(35) including COX-1. The data presented here for the complex membrane setup was previously published in that study. The exact compositions are given in Table S2. In brief, the membrane model contains an asymmetric distribution of the following major lipid groups: cholesterol (CHOL), phosphatidylcholine (PC), phosphatidylethanolamine (PE), and sphingomyelin (SM) placed in both leaflets; gangliosides exclusively in the upper leaflet, and charged lipids phosphatidylserine (PS), phosphatic acid (PA), phosphatidylinositol (PI) along with PI-phosphate, -bisphosphate, and -trisphosphate lipids (PIPs) placed exclusively in the lower leaflet. Complete details on the lipids used can be found on the MARTINI lipidome webpage (cgmartini.nl/index.php/force-field-parameters/lipids).

System equilibration was done using a gradual stepwise procedure, reducing the number and strength of position restraints on the proteins. Simulations were performed using a 20 fs time step. A target 310 K temperature was maintained with a velocity-rescaling thermostat(36), and a time constant for coupling of 1 ps. The Berendsen barostat(37) was applied semi-isotropically at 1 bar, a compressibility of 3 · 10^-4^ bar^-1^ and a relaxation constant of 5 ps. A small 1 kJ mol^-1^ nm^-2^ force constant on protein backbone beads was maintained during the simulation to prevent diffusion of the four protein copies and resulting proteinprotein interactions, which cannot be accurately sampled on a the time scale of the simulation. The system was simulated for 30 μs. Analysis of all CG systems, unless otherwise stated, was performed on the whole trajectory. All CG simulations were carried out using the GROMACS 4.6.x package(38).

### Atomistic Molecular Dynamics Simulations

For atomistic simulations, bilayer assembly, protein preprocessing and embedding into the bilayer, were done using the CHARMM-GUI webserver (39, 40). In addition to protein and lipids, the final system also contained TIP3P water(41) and 150 mM NaCl ions. The exact lipid composition of the atomistic systems is provided in Table S1. We used the CHARMM36m force-field(42) to describe the system and a 2 fs time step for integration. Particle Mesh Ewald(43) was used for long range electrostatics with a real space cutoff of 1.2 nm and 0.12 nm Fourier grid spacing. The same 1.2 nm cutoff was also used for van der Waals interactions. The Nosé-Hoover thermostat(44, 45) was used to maintain a target temperature of 310 K with a 1 ps coupling constant, applied separately to protein, membrane and solution components. We used the Parrinello-Rahman barostat(46) to keep the pressure at 1 bar, with a coupling constant of 5 ps and compressibility of 4.5 · 10^-5^ bar^-1^. The LINCS algorithm(47) was used to constrain bonds with hydrogen atoms. Bilayer composition includes simple lipid mixtures, composed of POPC lipids and either cholesterol (CHOL) or arachidonic acid (ARAN). Simulations that include large membrane are composed of only POPC lipids.

We performed several control simulations which differed in terms of their simulation parameters. Two of these systems were simulated using a surface tension of 5 and 30 *mN/m* applied in the x-y plane, respectively, and compressibility of 4.5 · 10^-5^ bar^-1^. To rule out any force-field dependency of our results one system was simulated using the GROMOS 54A7 force-field(48) and contained SPC water(49) instead. All atomistic simulations were carried out using GROMACS 2016.x package(50).

### Analysis

The GROMACS g_select tool was used to calculate the lipid count around proteins within a distance cutoff for the upper leaflet. Calculations for the lower leaflet were done using in-house scripts which used MDTraj(51) to process the trajectories, in addition to standard python libraries for scientific computing *(numpy* and *scipy)* and visualization *(matplotlib)*. 2D Density, thickness and curvature profiles calculated used an in-house developed program that has also been applied previously(35, 52) and explained in detail elsewhere(35).

Calculations of lipid contacts were done by considering both the total number of lipid contacts as well as their duration. When measuring the duration of contacts, we only consider the contact with the longest duration. These results were either plotted as a time series, as in Figure 2, or projected onto the surface of individual residues and colored via a color gradient, as in Figure 3.

**Figure 2.**
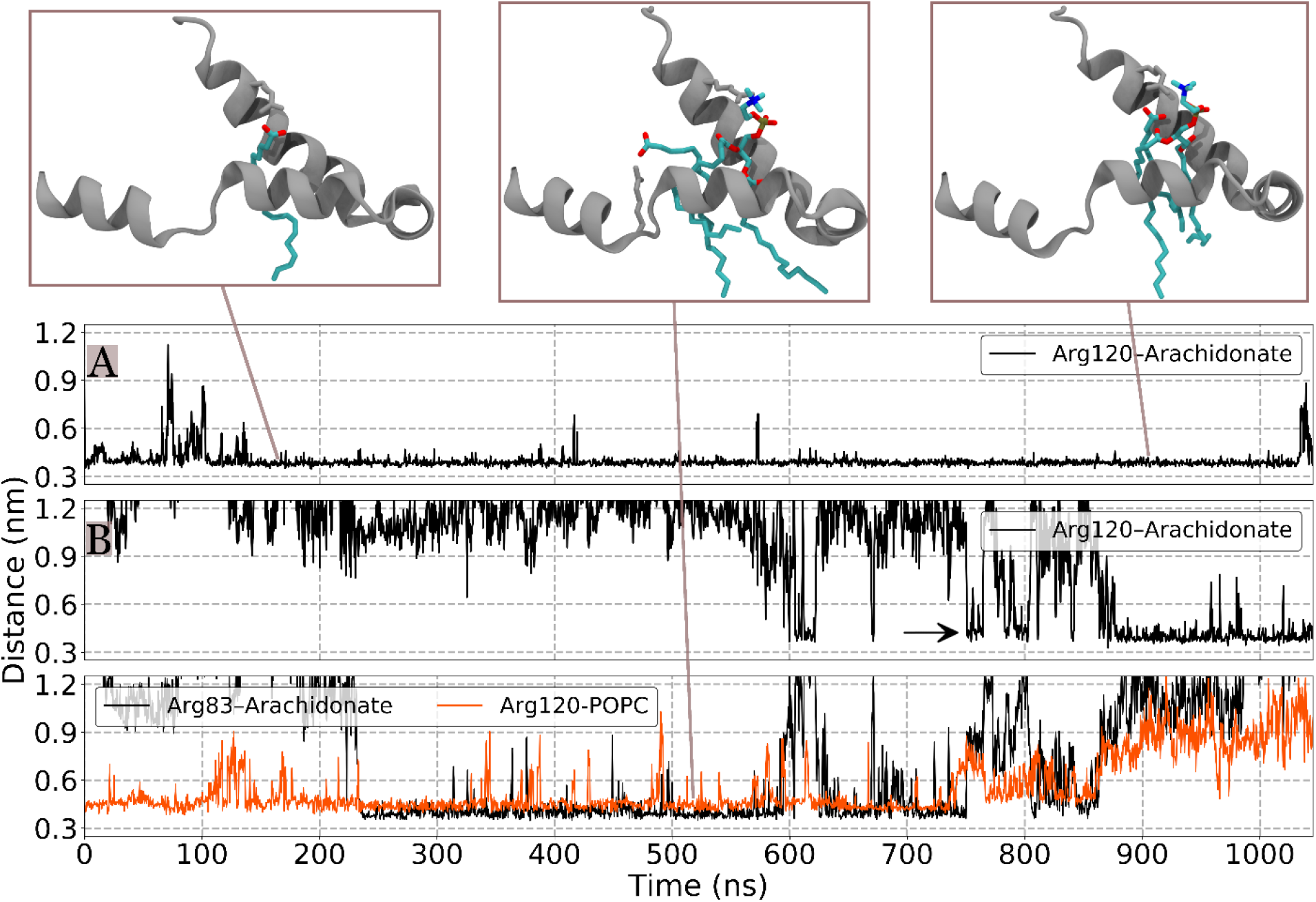
Arachidonic Acid (ARAN) binding to the MBD of COX-1 in AA simulations. Distance calculations between arachidonic acid carboxylate end and Arg-120 guanidino moiety for one monomer of COX-1(top line plot; denoted with A). For the other monomer (denoted with B) in addition to the same distance, we also show the distance between the phosphate headgroup of a bound POPC lipid to Arg-120. While the binding site is occupied, arachidonic acid interacts with Arg-83 instead (bottom line plot). The time when arachidonic acid starts interacting with Arg-120 is marked with a black arrow.

To show bilayer deformations as a result of POPC lipid interactions with basic residues on the surface of cyclooxygenases, we calculated distances from lipid P atoms and COX-1 center of mass. These calculations were done for three mutually exclusive categories, meaning that a POPC lipid can at most belong to one of these categories. They are: POPC lipids in direct contact with arginine or lysine residues, POPC lipids in close contact with the whole protein (excluding those interacting with arginine and lysine residues), and other POPC lipids. To define what constitutes a ‘direct contact’ we used three different distance cutoffs: 0.4, 0.5 and 0.6 nm. The results presented in Figure 6C are derived from the system with a small surface tension applied. This was done to decouple bilayer undulations, which is a property of every biological membrane, from perturbations to the membrane, such as positive curvature, caused by the presence of COX-1. The application of surface tension will eliminate bilayer undulations and allow us to measure the effect of COX-1 on the curvature of the membrane.

Visualizations are done using VMD(53) and NGL Viewer(54, 55).

## Results

### Characterization of arachidonic acid binding to COX-1

The endogenous substrate of COX-1 enzymes is arachidonic acid which is released by phospholipase A_2_ – mediated hydrolysis of glycerophospholipids(56). Computational studies of COX-1 interactions with arachidonate focus on interactions of the lipid already bound at the cyclooxygenase site(6, 7, 21, 22) of the enzyme with residues surrounding it. The way the lipid enters the active site, however, has received much less attention. In atomistic simulations of COX-1 embedded in a membrane model containing arachidonate, we observe the entrance and specific binding of the latter into the hydrophobic channel of the MBD (Figure 2). Interactions with residues lining up helices A-D of the MBD lead to arachidonate being pulled out of the membrane plane and its insertion into the hydrophobic cavity. This configuration with the lipid bound inside the MBD hydrophobic core is highly stable and maintained throughout the simulation. We identify two types of interactions that are responsible for this binding: charge-charge interaction of the carboxylate end of arachidonate and guanidinium moiety of Arg-120, as well as the hydrophobic interaction between the arachidonate acyl chain and hydrophobic residues lining up helices B and D of the MBD.

To measure the binding of arachidonate to the hydrophobic channel of COX-1 we performed distance calculations between the carboxylate headgroup of the lipid and the guanidinium moiety of Arg-120. We chose Arg-120 as a reference residue for these calculations throughout this study for the following reasons: *(i)* Arg-120 forms the entrance to the COX-1 cyclooxygenase site and as such delineates its entry point, *(ii)* it interacts with and stabilizes the bound lipid headgroup, *(iii)* it is of great functional importance to the activity of COX-1(57, 58) and *(iv)* the relative alignment of the distance vector connecting the lipid to Arg-120 is close to the bilayer normal vector, and as such it simultaneously allows us to measure the degree of lipid insertion into the channel. These results are highlighted in Figure 2B. These distance calculations reveal that once arachidonate binds to the hydrophobic channel of COX-1, it is maintained there for the whole duration of the trajectory with minimal fluctuation, which is highly indicative of specific binding.

COX-1 is a homodimer, each monomer containing a catalytic site and hence, a hydrophobic channel for lipid entrance. We observe the immediate binding of arachidonate on only one of the monomers. In the other monomer, however, we observe the binding of a POPC lipid instead. This binding – which has been reported before in the MD literature for DMPC(23) – is more variable and therefore less stable, hinting at the interaction being nonspecific and in the long run more likely to dissociate. Nevertheless, in our simulations, while bound, POPC clearly blocks arachidonate from entering the site and disallows binding. Instead, it is kept at the ‘front door’, where it interacts with Arg-83. During the course of arachidonate interactions with Arg-83, we see the lipid coming in close contact with Arg-120 several times (Figure 2, black arrow). After close to 900 ns, we observe the displacement of the POPC lipid by arachidonate but do not observe it completely removing POPC out of the MBD, which we expect to happen given more sampling. The MBD expands to accommodate both lipids at this site: POPC interacting with mainly helix B and arachidonate interacting with helix B and D residues.

The binding poses we observe for both lipids that bind to COX-1, include the lipid “extracted” from the membrane and reaching heights higher than the average membrane plane. Arachidonate for instance, when bound, reaches a height of 1.98 ± 0.17 nm compared to 1.65 ± 0.05 nm for all other arachidonate lipids combined (calculated over the last few frames of the trajectory, distance is relative to membrane center). Such a lipid configuration is stable and thermodynamically viable as a result of the aforementioned charged-based interactions of the lipid headgroup with Arg-120 and hydrophobic interactions of mainly Val-116 and Val-129, but also other hydrophobic residues from helices B and D of the MBD. In a similar simulation with equal length but using a protonated (uncharged) arachidonic acid, we still see it binging to the MBD and interacting with the same residues, highlighting thus the importance of hydrophobic interactions in maintaining the binding.

### Coarse-grained MD simulations reveal a complex interplay of COX-1 enzymes with lipids

The occupancy of the cavity within the MBD of COX-1 by POPC lipids, closes the pathway used by arachidonate to access the enzyme’s active site. From these results, however, it is unclear if this binding is specific to POPC lipids or perhaps shared by other lipids as well. To test this, we conducted CG MD simulations of the enzyme embedded in several model bilayers. The construction of these membrane models aimed at reproducing the unique lipid composition of ER membranes and include several lipid species. Specifically, we employed low concentrations of cholesterol which in ER membranes is reported to be in the range 5-8% (30–32) and a large ratio of PC lipids (Table S1). These systems also contain very small amount of sphingolipids to match the low concentration of sphingo- and other complex lipids in ER membranes(59, 60). Figure 3A shows 2D density maps calculated separately for the upper and lower leaflet and highlighting the preferential localization of lipids in one of these systems (Figure S2 shows the same calculations for all systems combined). Owing to their monotopic binding to the upper leaflet of the bilayer, COX-1 interactions with lipids are defined predominantly by lipid interactions with their membrane anchoring domain.

**Figure 3.**
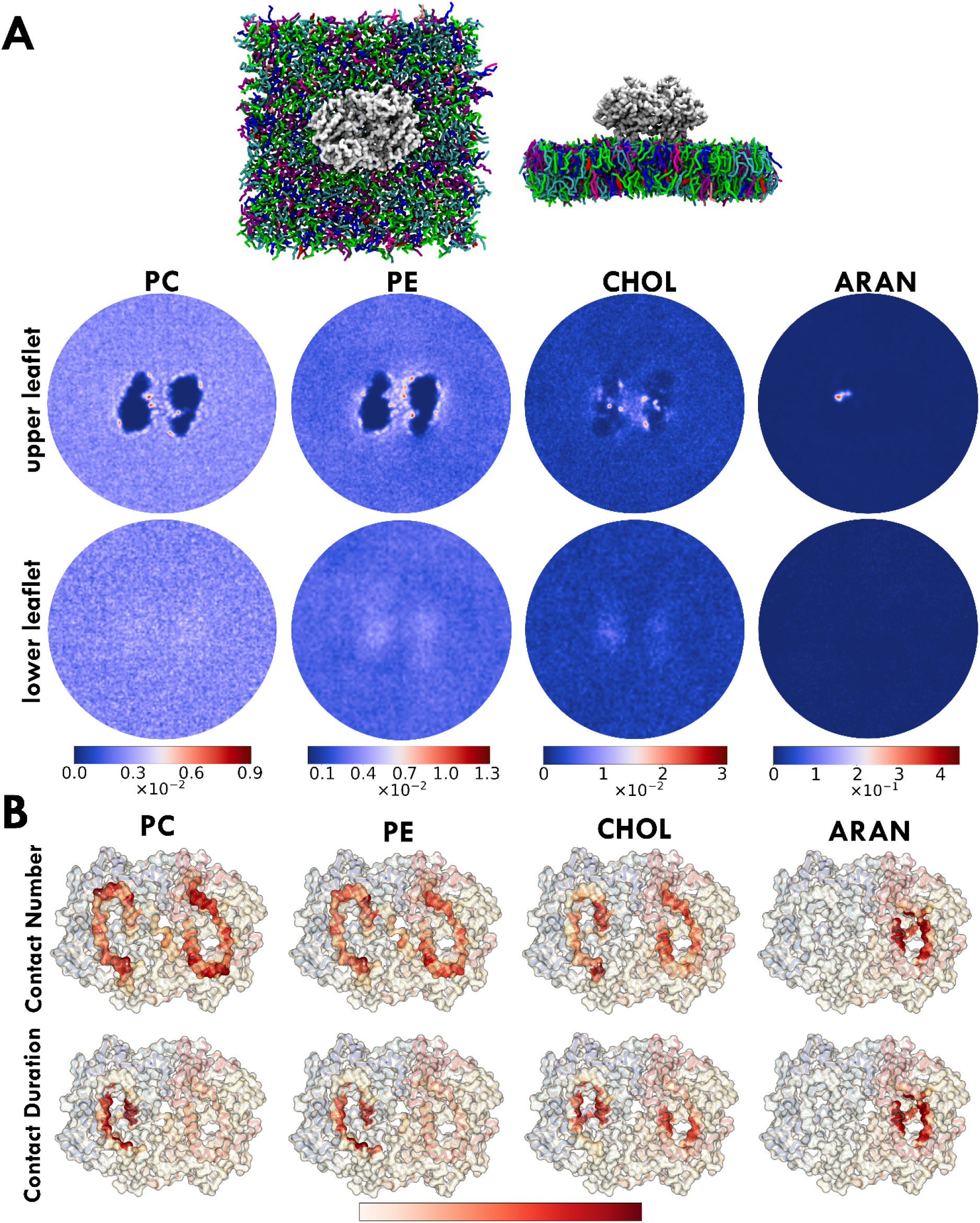
COX-1 interaction with lipids in ER-like membranes. A. Top and front view of the simulated system with each lipid type colored using a different color, along with density maps calculated separately for the upper and lower leaflet for PC, PE, CHOL and ARAN. B. Contact heatmaps between COX-1 and the same lipid groups calculated as the total number of contacts or their respective duration, using a white-red color gradient.

The MBD inserts well into the width of the upper bilayer leaflet and interacts with surrounding lipids. Most of these interactions are short-lived and transient; we do, however, also observe several interactions that persist for much of the simulation time. Such interactions hint at the possibility of specific lipid-protein interactions, which are interactions that may serve a functional or modulatory role in the activity of proteins. Figure 3A shows density maps for the following lipid groups found on the upper leaflet of the membrane: CHOL, PC, PE, and arachidonic acid (ARAN). The upper leaflet also contains SM lipids, but since they do not show any notable interaction with the enzyme, they are omitted from the figure. We see that each of these lipid types forms highly localized interactions with COX-1. Specifically, we note that the majority of these interactions are located within the interface formed by the MBD of each monomer, and in particular we observe that each of these lipids is capable of interacting with the hydrophobic channel inside the MBDs. To further highlight the latter, Figure 3B shows contact heatmaps between COX-1 and each of the above lipids measured as either their total number of contacts or their respective duration of interaction. Calculations are done on a per-residue basis and the resulting heatmaps are projected onto the surface of the enzyme. The heatmap formed by the total number of contacts confirms that the majority of interactions include the MBDs of COX-1. When we look at the longevity of these interactions instead, however, we see that only those interactions that involve the hydrophobic channel of the enzyme are left, showing clearly that only lipid interactions with this site of the enzyme are maintained for prolonged durations of the trajectory.

When comparing the binding of different non-arachidonate lipids against each other in terms of their consistency and specificity of binding, despite its low concentration in our model membranes, we find that in the majority of cases, cholesterol is the dominant lipid species interacting with COX-1 and occupying the MBD cavity. We observe cholesterol binding even in setups where its content is very low (3% of total lipids) or arachidonate is one of the lipid components (Figure S2). Therefore, to highlight cholesterol binding we measured distance calculations of cholesterol bound at this site from a CG setup involving a complex membrane model composed of 63 different lipid types (but lacking arachidonate) wherein four copies of COX-1 have been embedded and that has been simulated for 30 μs. Figure 4 shows data for one of these copies and Figure S1 for the rest. These results indicate that despite the presence of many other lipid species in the system, we mainly see cholesterol occupying the hydrophobic channel of COX-1 and forming strong interactions that are maintained from a few and up to 13 μs of simulation time.

**Figure 4.**
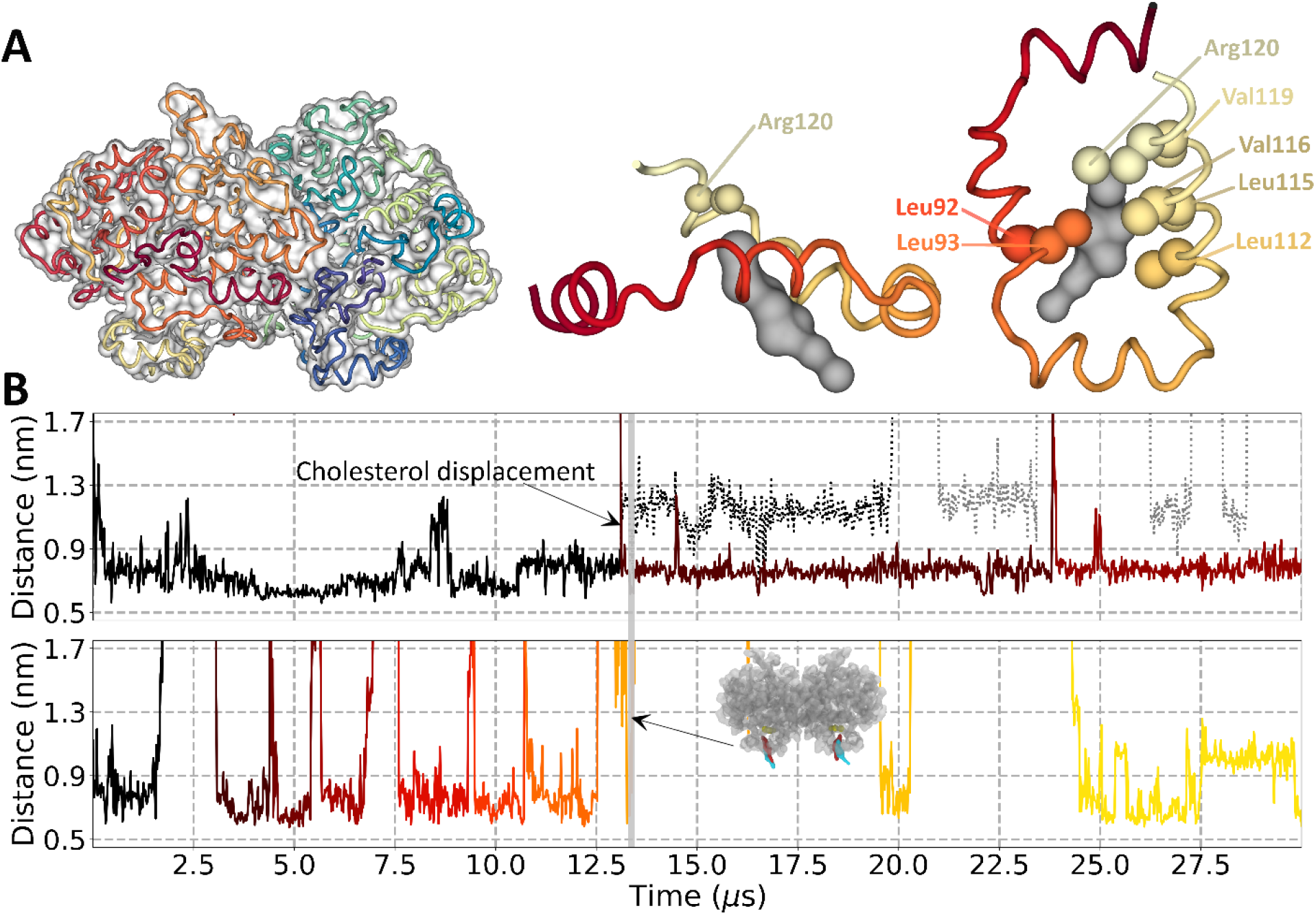
Cholesterol binding to the MBD of COX-1 in CG simulations. **A.** Overview of the binding site inside the MBD. On the left the backbone of the coarse-grained protein is shown along with a transparent surface representation. The two panels on the right highlight the interaction of cholesterol (gray surface) with helices A-D of the MBD. Residues that interact most with cholesterol as shown as sphere representation. (coloring and viewing angles are similar to Figure 1). **B.** Arg-120 – cholesterol distance as measured by the MARTINI representation of the arginine guanidinium and cholesterol hydroxyl moieties (ROH-SC2 beads) measured separately for each monomer. The same distance for the second cholesterol molecule that is sometimes bound at roughly the same site is drawn with a greyed out dotted line.

Figure 4A provides a close view of the binding site within the hydrophobic channel of the protein along with residues involved in the binding process. The binding site is formed by the hydrophobic sidechains of leucines 92, 93, 112, and 115, as well as Val-116, and Val-119, which are positioned at opposing sides of the binding site (helices B and D) and form a tight interaction network with the hydrophobic ring structure of cholesterol. The hydrophilic headgroup of cholesterol, on the other hand, interacts with Arg-120, an essential residue for the activity of COX-1(57, 58). Figure 4B shows distance calculations for both COX-1 monomers between the cholesterol headgroup (ROH bead in the MARTINI model) and the guanidino moiety of Arg-120 (SC2 bead). This measurement was done to match the distance calculation shown in Figure 2 for arachidonate.

In our simulation setup we have four enzyme copies, each composed of two monomers. Hence, the data that we show in Figure 4 (and Figure S1) correspond to 8 total binding sites. Collectively, we see several cholesterol binding and unbinding events, associated with varying levels of close contact durations, but generally no less than 2 μs. We also observe that the COX-1 hydrophobic channel is sometimes occupied by a second cholesterol molecule (Figure 4B, dotted line), with a much sparser frequency and lower longevity of binding. While the MBD certainly seems capable of accommodating two cholesterol molecules, this is not a stable configuration. This could also be a consequence of the presence of an elastic network on CG proteins, which would prohibit the MBD from expanding to accommodate both lipids (similar to what we observed for arachidonate).

The combined results from CG simulations confirm the binding of arachidonate similar to what we observe in atomistic simulations, and further show that other lipids – mainly, cholesterol, but also PC and PE lipids – can also bind at the same interaction site as the endogenous ligand of COX-1. The biological implications of these findings are twofold: *(i)* the hydrophobic channel of COX-1 is usually in an occupied state and not empty, and therefore *(ii)* the dissociation of the bound lipid at this site has to precede arachidonate binding. The former relates to the most populated state adopted by COX-1 in ER membranes, whereas the latter hints at a possible importance of membrane lipids in affecting the kinetic profile of the enzyme. This is because the ability of a lipid to occupy a space, the accessibility of which is required by the native ligand for the biological activity of the protein, underscores a potential regulatory activity of the lipid in the activity of the protein.

### Atomistic simulations of cholesterol interactions with COX-1

To further investigate cholesterol binding to COX-1, we carried out simulations at the all-atom level of detail using simpler membrane models (POPC:CHOL). In simulations where the level of cholesterol is low (~15% of bilayer lipids), we do observe some prolonged interactions of COX-1 with cholesterol, but we do not observe any binding to the hydrophobic channel. While it may be the case the atomistic simulations would result in different interactions compared to the coarse-grained simulations, we believe that the more limited sampling in the former is a far bigger issue. To account for this, we carried out the same simulations in setups with a larger cholesterol content, and indeed we do observe cholesterol binding to COX-1, at the same site as in the CG simulations (Figure 5). The binding site itself, as well as the residues that are responsible in maintaining the binding are the same as what we saw from CG simulations (see the COX-1 surface representation comparison between all-atom and coarse-grained results in Figure 5C).

**Figure 5.**
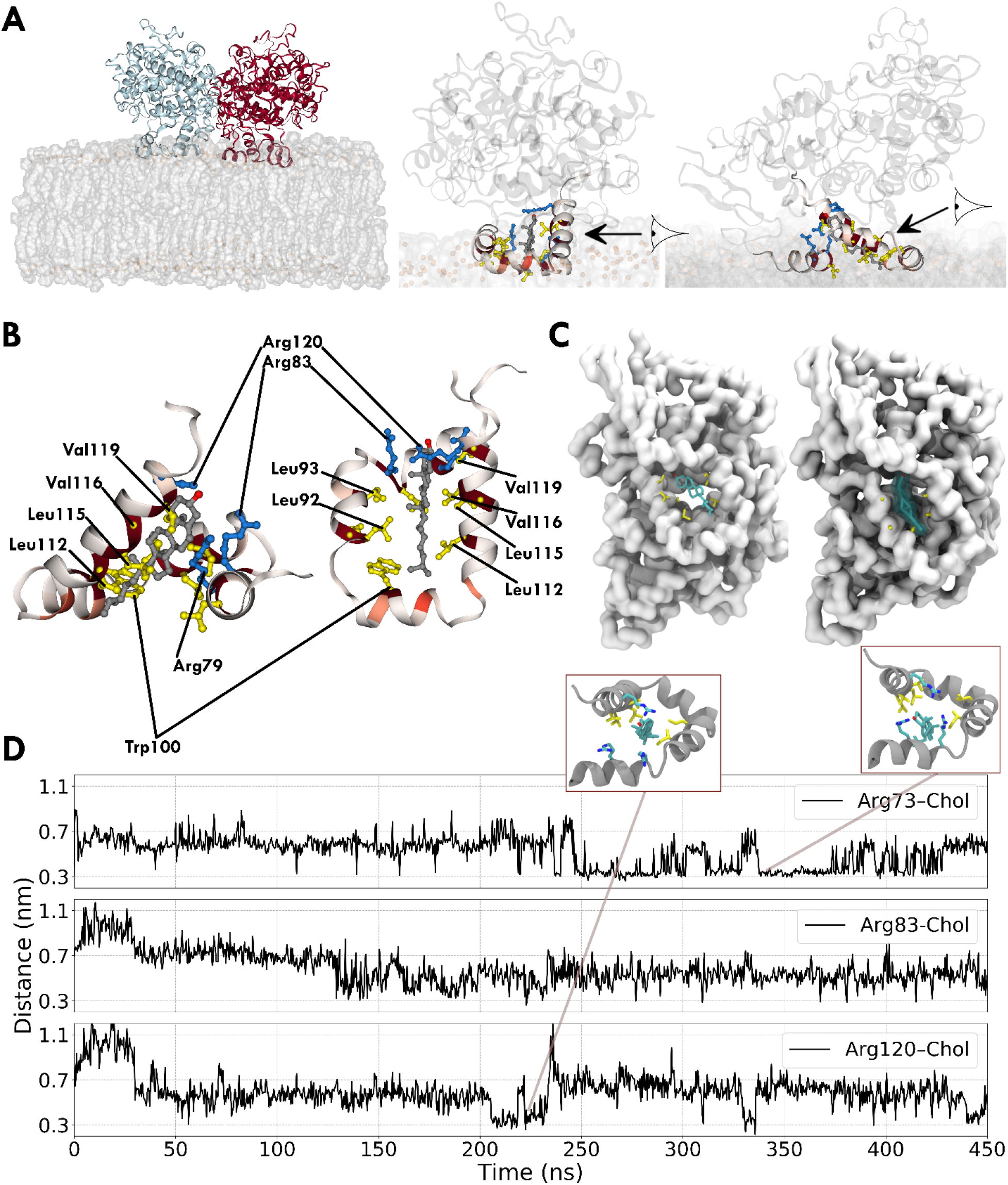
Cholesterol binding to the MBD of COX-1 in AA simulations. **A.** Overview of the simulated system anchored to the membrane viewed in full (left) and the monomer bound to cholesterol (middle and right). B. Closed-up view of the bound cholesterol. Hydrophobic residues lining up the interface are shown in yellow. Arginine residues are drawn in blue. The coloring of the MBD uses the same color gradient as Figure 3 based on the number of interactions with cholesterol that are formed. C. Comparison of the binding site in the atomistic (left) and CG simulation (right). D. Distance calculations between the hydroxyl oxygen of cholesterol and the guanidinium carbon of three arginine residues that stabilize the polar headgroup of cholesterol.

The rough surface (β-face) of cholesterol faces the inside of the protein, with the smoother (α-face) directed towards the membrane. Similar to the CG simulations, a network of hydrophobic residues (Leu-92, Leu-93, Leu-112, Leu-115, Val-116, and Val-119) lining the MBD stabilize the ring structure of cholesterol. Somewhat differently from the CG simulations, however, three adjacent arginine residues interact with its hydroxyl headgroup. Figure 5B shows distance calculations from these three arginine residues to the cholesterol hydroxyl headgroup, calculated to make comparison with the CG data easy. Binding occurs at the start of the simulation and is maintained throughout. Furthermore, all three arginine residues form contacts with cholesterol, with its hydroxyl headgroup placed equidistantly between them with shorter intervals where cholesterol interacts predominantly with either Arg-73 or Arg-120. For Arg-73 binding to occur, helix A bends towards the other helices. In another similar simulation, however, we observe cholesterol binding without the bending of helix A and the involvement of Arg-73, as such interactions with Arg-73 are not necessary for cholesterol binding.

In the setup with ~15% CHOL content, where we did not observe cholesterol binding, instead we see a POPC lipid insert and interact with residues at roughly the same site as arachidonate and cholesterol (Figure S3). In contrast to them, however, POPC binding appears to involve largely electrostatic interactions. Even though the 2-oleyl tail interacts with Trp100 of helix C, the bulk of the interaction stability comes from interactions between the lipid phosphate and Arg-83 guanidino moieties (as is evident in Figure S3, where when this interaction breaks and the choline headgroup starts interacting with the very vertically placed Glu-493, the overall positioning of POPC at this site becomes less well-defined). This interaction with POPC inside the MBD hydrophobic channel in the setup with low cholesterol content may explain why we did not observe cholesterol binding there, since for it to occur we would have to sample the dissociation of the bound POPC lipid first. That is, the prior binding of POPC increases the sampling time required to observe the binding of cholesterol (note that this does not convey any information about their relative strength of binding).

Overall, while binding of cholesterol to COX-1 is specific, the binding site itself does not seem to be exclusive to only cholesterol (as also noted above from CG simulations). The overall conclusion of atomistic simulations seems to be a strong support for cholesterol binding, with a side note that POPC (and likely other lipids) as well can access the same interaction site.

### COX-1 induces a positive curvature on the surrounding lipid environment

Measurements of membrane physical properties such as membrane thickness, curvature, shear stress, strain and elasticity profiles show that proteins in addition to forming specific interactions with individual lipids, can also cause perturbations to the local membrane environment(18). The latter is the result of nonspecific lipid-protein interactions. These “bulk-lipid” effects have been studied extensively but generally are less well understood compared to the specific interaction with lipids. In our simulations, regardless of the setup, lipid composition or resolution of the models used, we consistently observe COX-1 exhibiting a marked effect on the surrounding membrane environment.

Despite their localization on the outer leaflet of (mainly) ER membranes, in all our setups we see COX-1 induce structural changes to the membrane and affect both leaflets. The immediately observable visual manifestation of these perturbations is the creation of a positive curvature around the enzyme (Figure 6). Calculations of the mean curvature profiles for the CG systems highlight the ability of COX-1 to affect the curvature of the upper and lower leaflet (Figure 6D). This effect is also easily observable when calculating the average thickness of the membrane model during the simulation, where for the same system, we see a higher average thickness at the enzyme embedding location compared to the surrounding environment. It is clear that this effect is not a result of specific lipid-protein interactions. Rather, it is a consequence of COX-1 nonspecific interactions with membrane lipids. In Figure 3A and, more prominently Figure 6B, we notice that in addition to lipids interacting with residues inside the hydrophobic channel of the enzyme, they also preferentially localize at the interface between its MBDs, forming many smaller and less well-defined interaction sites. This is in particular the case for phospholipids.

**Figure 6.**
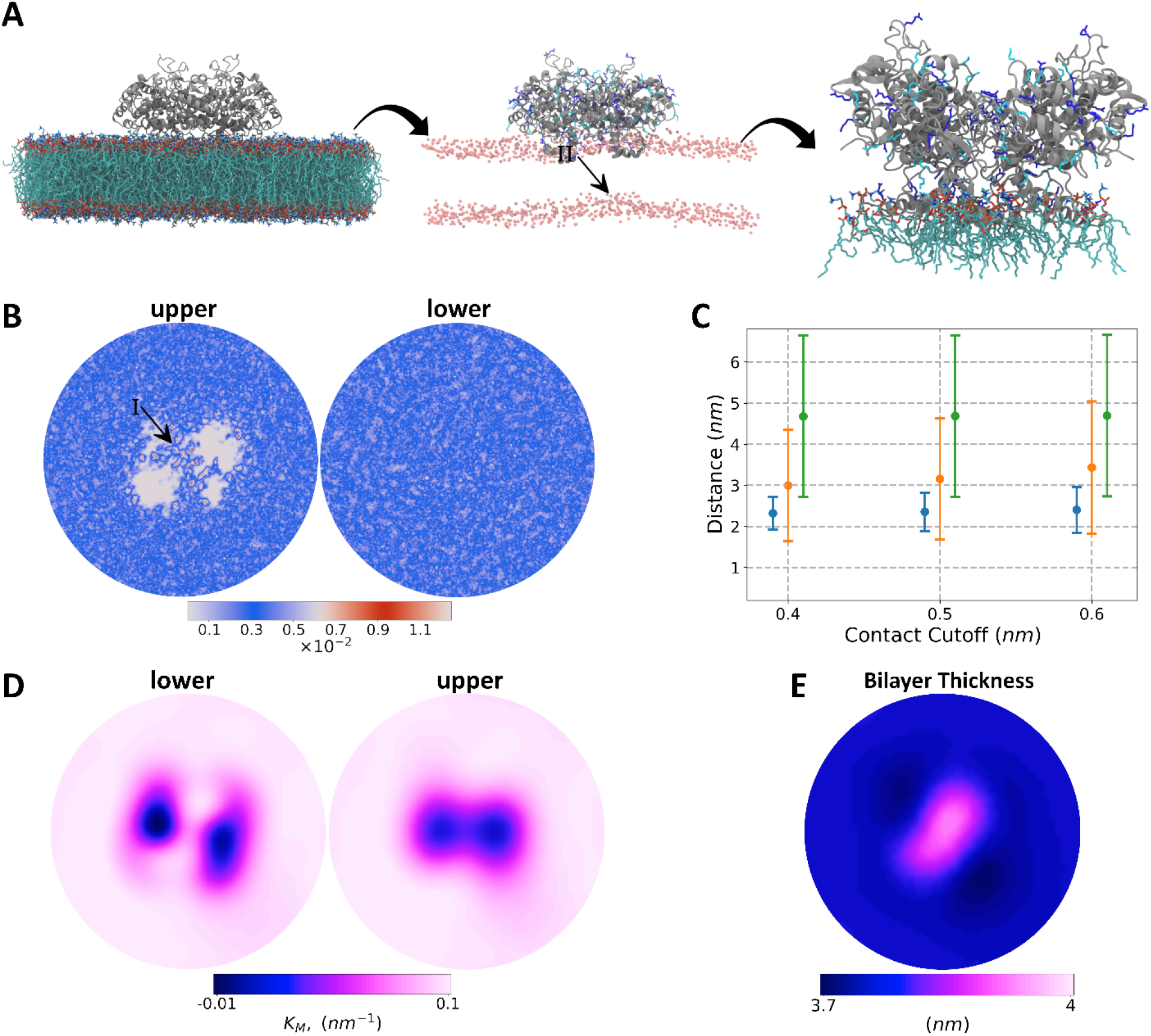
Curvature inducing property of COX-1. **A.** Schematic view of bilayer perturbations as a result of COX-1 interactions with POPC lipids. The middle panel shows the changes to the lower leaflet by only visualizing the POPC phosphorus atoms (arrow II). Positively charged residues are colored with dark and light blue for arginine and lysine, respectively. **B.** 2D density profiles for the lower and upper leaflet highlighting the increased localization of POPC lipids underneath the enzyme (arrow I). Please note the colorbar follows a ‘cyclic’ color gradient (see reference (61) for more information) for easy identification of preferential localization sites. **C.** Distance calculations between COX-1 center of mass (COM) and P atom of POPC lipids, from three mutually exclusive groups: POPC lipids interacting with arginine or lysine residues (blue), POPC lipids interacting with residues other than arginine and lysine, and all other POPC lipids. The definition of contact is done using three distance cutoffs: 0.4, 0.5, and 0.6 nm. **D** and **E.** Visualization of the mean curvature and membrane thickness, respectively.

To test if these non-specific interactions with membrane phospholipids are the mechanism by which COX-1 enzymes perturb the local environment, we carried out all-atom simulations of COX-1 embedded in a pure POPC membrane. These simulations employ a large membrane setup, which is necessary to distinguish between local lipids and bulk bilayer lipids. Figure 6A shows snapshots of one of these systems visually highlighting the curvature the upper-leaflet embedded COX-1 induces on the overall membrane (*arrow II*). The resulting density profiles of POPC lipids during the simulation (Figure 6B) confirm their increased localization at the interface between each monomer’s MBD (*arrow I*). COX-1 contain many surface-exposed positively charged residues, with several facing the membrane where they interact with local lipids. The interaction of phospholipids with these residues ultimately leads to the creation of the positive curvature we observe in our simulations. To show this, we separated POPC lipids into three mutually exclusive groups: POPC lipids interacting with lysine or arginine residues (blue), interacting with all other COX-1 residues (orange), and all other POPC lipids (green) and calculated their average distance from the center of mass of COX-1 (Figure 6C).

The cutoff for what constitutes a contact/interaction is done for three different distances: 0.4, 0.5 and 0.6 nm. The result shows that POPC lipids interacting with positively charged residues are on average closer to the center of mass of the protein then other POPC lipids, revealing thus a likely mechanism for the capability of COX-1 to induce a positive curvature. First, the MBDs act as physical barriers that separate lipids into protein-interacting, low diffusion lipids and freely moving, high diffusion lipids. Since proteins in general affect the diffusion rates of lipids, there are two additional factors that play a key role for cyclooxygenases. The outer surface of COX-1 is characterized by a plethora of positively charged residues which form charge-charge interactions with lipids containing negatively charged chemical groups (predominantly phospholipids) resulting in a height difference between lipids in-between the MBDs and surrounding lipids. And lastly, the partial insertion of COX-1 into the membrane leaves the lower leaflet without any supporting protein interactions. The combined result is the induction of a positive curvature on the local membrane environment, which is persistent even in simulations employing surface tension, or other control parameters (see methods). In contrast to the upper leaflet, calculations of lipid order parameters and number densities reveal no changes to their values for the lower leaflet (Figure S4-S6), further supporting our claim that the curvature creation is driven purely by COX-1 interactions with lipid in the upper leaflet and the enzyme does not affect the lipid composition of the lower leaflet.

## Discussion

The understanding of the interplay between membrane embedded proteins and their surrounding lipid environment has become an important part of protein function and activity studies. MD simulations have revealed the detailed lipid interaction profile of many proteins (including GPCRs(52), ion channels(62), and many others(18)).

The endoplasmic reticulum is the main organelle for lipid synthesis accounting for the majority of phospholipids (e.g. PC, PE, PS etc. lipids)(63) and cholesterol(31). They are, however, quite low in cholesterol content itself, resulting in a loosely packed configuration of lipids which fits their role in transporting synthesized lipids to other organelles. Cyclooxygenases are bound monotopically to the outer leaflet of ER membranes. In addition to their evolutionary adaptation to function in such a highly fluid membrane environment, we find that COX-1 is itself capable of inducing a positive curvature. The effect is persistent regardless of the lipid composition of the membrane model and simulation setup used. The observation of this property in MD simulations, to our knowledge, was done independently by Wan et al.(24) using atomistic simulation and Balali-Mood et al.(27) employing a coarse-grained resolution. In the former, the authors observed the curvature of both cyclooxygenase isoforms during 25ns runs, whereas in the latter study, the authors show that monotopic membrane proteins with a deep insertion into the membrane cause bilayer perturbations similar to those presented here (although the degree of the curvature was much more pronounced in their work, which may be due to their use of a nascent version of the MARTINI model(64)).

Through a combination of membrane curvature and thickness profile calculations coupled with measurements of order parameters and density maps from both atomistic and coarse-grained resolution, we are able to provide a mechanistic explanation for these perturbations. Nonspecific yet prolonged interactions of surface-exposed positively charged residues with membrane phospholipids located at the interface between the MBDs of the enzyme provide the driving force leading to the creation of the observed curvature around COX-1. The non-specificity of these interactions stems from the fact that in our simulations they are driven by charge-charge interactions of the lipid phosphate headgroup with arginine and lysine residues. In our atomistic simulations we chose POPC to model the presence of phospholipids. Since ER membranes are composed of around 85% by phospholipids(30) and the COX-1 induced curvature is charge-driven, we are confident that our results are invariant to the details of the used membrane models and simulation parameters.

In contrast to these local and non-specific interactions of COX-1 with phospholipids that covers the entire interface between the MBDs, we observe specific interactions with membrane lipids at only one interaction site per monomer. This interaction site is located inside the hydrophobic channel created by the four helices of the MBD. In our CG simulations, we see PC and PE lipids, but not SM lipids, interact with residues lining up the interface of the channel and occupying it for prolonged durations of time. In addition to these phospholipids, we also observe the binding of cholesterol at the same site. Altogether, the MBD is able to accommodate a variety of phospholipids and cholesterol (in addition to its endogenous ligand), revealing a picture of the enzyme which shows a lipid bound at its MBDs for the majority of the time. In terms of frequency of binding as well as strength of binding – measured here as the maximum duration a lipid is in contact with the same (set of) residue(s) – we find cholesterol and arachidonate to be the main lipids in occupying this interaction site.

The binding of arachidonate, in particular, is marked by a high stability and small fluctuations. In one monomer, arachidonate binds directly to Arg-120 and is maintained there for the whole simulation. In the other monomer, however, a POPC lipid binds first and occupies the site, disallowing arachidonate from doing the same. POPC binding, however, is less stable and as the simulation progresses it is eventually displaced by arachidonate (Figure 2). In this case, when the MBD is occupied by another lipid, arachidonate binding follows a different pathway. We calculated the number of contacts that this arachidonate forms during the simulation (Figure 7A) and see that 4 out of 5 most prominent interactions are with arginine residues. These arginine residues are located in the EGF domain of the enzyme (Arg-61) and along the length of the MBD and form the pathway for arachidonate binding (Figure 7B). Charge-charge interactions between arachidonate carboxylate and arginine guanidino atoms allow the former to bind temporarily to the latter. Due to the lack of any stabilizing hydrophobic residues, these interactions are not stable, and only serve to allow arachidonate to sample interactions with other residues in the vicinity. This way, these four arginine residues act as “connecting bridges” whereby arachidonate “jumps” from one to the other and ultimately binds to Arg-120, from where it enters the active site of the enzyme (we do not observe this process in our simulation). The main arginine residue in this pathway is Arg-83, which forms the most interactions with arachidonate and is responsible for keeping it close to the binding site when it is occupied by another lipid and enabling its interaction with Arg-120. Upon binding to Arg-120, arachidonate is stabilized via hydrophobic interactions with valines 116 and 119, replaces the bound POPC lipid, and is maintained there for the rest of the simulation.

**Figure 7.**
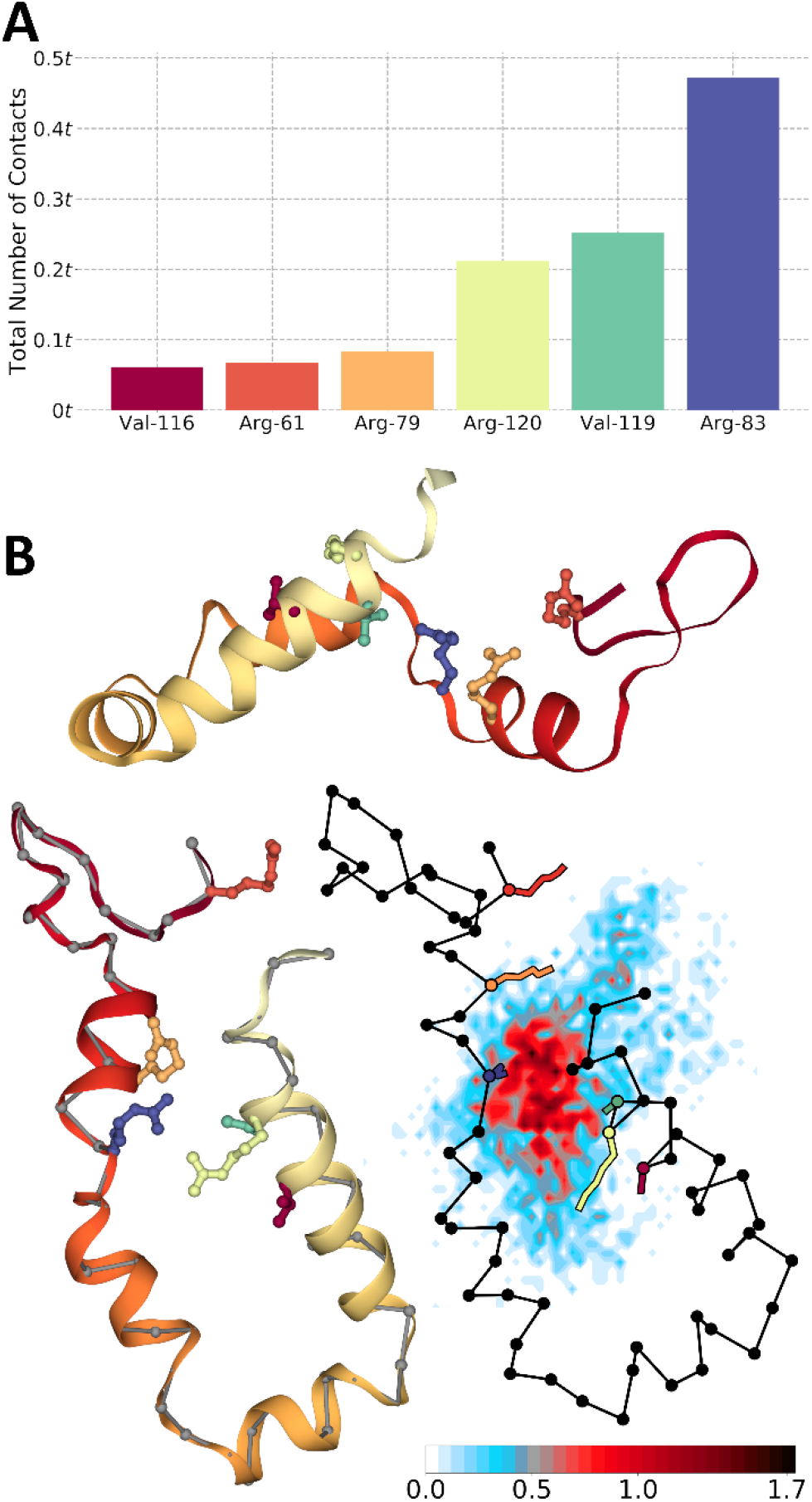
Arachidonic acid binding pathway. **A.** The sum of all contacts formed between arachidonic acid and COX-1 residues during the whole simulation. The six residues with the highest number of contacts are shown. **B.** Side and top view of the MBD with arginine residues forming the pathway shown with a stick representation along with the density map of arachidonic acid during the simulation showing its pathway. On top of the density we overlaid the position of MBD residues (colored in black) and the six residues with the most contacts. Positions are averages during the simulation, and residue colors are consistent between subplots. t is the simulation time.

The other lipid that we observe to interact specifically with COX-1 is cholesterol. Its binding, similar to that of arachidonate and phospholipids, includes the hydrophobic channel inside the MBD. We observe it consistently and independently in both CG and AA simulations involving largely interactions with the same residues. Binding of cholesterol to COX-1 is achieved by interactions of cholesterol headgroup atoms (ROH bead in MARTINI, hydroxyl in all-atom simulations) with arginine residues of helix B and D (Arg-83 and Arg-120). The core structure of cholesterol forms interactions with several hydrophobic residues lining up the interior of the MBD. In our simulations, cholesterol binding is observed more frequently and with longer duration compared to the binding of phospholipids, however, future studies looking at the free-energy landscape of these interactions are required to provide a more definitive characterization of their relative binding strengths. In addition, longer simulations are necessary to probe arachidonate insertion into the active site of COX-1.

Only one of the COX-1 monomers is catalytically active at a given time (Ecat) and it is modulated allosterically by its partner monomer (Eallo). The cooperative cross-talk between the subunits is an important determinant of their function and is heavily dependent on ligand binding. Palmitic acid, for instance, has an opposite and reverse allosteric effect for the cyclooxygenase isoforms, activating COX-1 by a factor of 2 and inhibiting COX-1 by ½(65). Other nonsubstrate fatty acids (e.g. stearic and oleic acid) have the same effect on COX-1. Measurements of binding kinetics for these fatty acids indicate that their inhibition of COX-1 may not be due to competition with arachidonic acid for Ecat(15). These studies reveal a complex and isoform-dependent interplay between fatty acids and cyclooxygenases. In a recent review, Smith and Malkowski(15) conclude that cyclooxygenase activity is both pharmacologically and physiologically dependent on the fatty acid content and flavor of the membrane environment. MD simulation results presented here broaden our perspective on COX-1 binding to ER membranes. First and foremost, we identify a previously unexplored and undefined pathway for arachidonic acid entrance into the COX-1 active site. Further, we show that the cavity within the MBD is occupied by membrane lipid for the majority of the time, which indiscriminately binds to fatty acids, cholesterol and phospholipids. Regarding the latter, we do not observe a big difference between different phospholipids binding. Binding of phospholipids without an obvious preference between them has been reported before in the MD literature for ABC transporters(66). Future calculations of the free-energy landscape of these binding events would reveal a better picture of COX-1 preference for phospholipids. Our results, however, agree with experiments of reconstituted COX-2 in nanodisks containing different phospholipids which showed no effect of changing phospholipids on the activity and inhibition of the enzyme(67). Considering the often-different ligandbinding outcomes for cyclooxygenase isoforms, however, it is unclear if these results hold for COX-1 as well.

MD simulations, in addition to providing insight into the interplay between molecular components, should also guide experimental work. The discovery that the COX-1 hydrophobic channel harbors specificity of binding to nonsubstrate fatty acids(68, 69) and, in our work, non-fatty acid lipids opens the possibility of membrane lipids playing a functional role in the activity of these enzymes. This could be in particular the case for cholesterol, which is found in small concentrations in ER membranes, and thus may provide cells with a regulatory mechanism. The results presented here, indicate that the kinetic profile of COX-1 should be dependent on the relative ratios of membrane lipid components – something that has already been shown in the case of fatty acids(16, 69) Additionally, we identify Arg-83 and hydrophobic residues Val-116 and Val-119 as important to ensure proper binding of arachidonic acid, especially if the MBD is already bound to a different lipid. Experimental techniques such as site-directed mutagenesis could be used to test the validity of these findings. Phe-205 mutants, for example, have shown a several-fold decrease in enzyme (COX-2) efficiency compared to the wild type(58). Single (R120A) and double (R120A/G533A) mutants of Arg-120 also affect the efficiency of COX-2, with the former increasing the Km value 3.4-fold without affecting the oxygenation of arachidonic acid, and the latter completely losing the cyclooxygenase activity(58). Mutants of Arg-83, as well as Val-116 and Val-119, could display a similar increase in Km (albeit perhaps to a lesser extent). Comparing the sequences of ovine, murine and human cyclooxygenases, similar to Smith et al.(9), we see that Arg-61, Arg-120 and Val-116 are conserved throughout, with Arg-79 and Arg-83 being conserved for COX-1 and replaced by a lysine for COX-2, which likely would serve the same function in guiding arachidonic acid via charge-charge interactions. In contrast, Val-119 is conserved among COX-1 but replaced with a serine in COX-2 enzymes.

## Acknowledgements

This work was supported by the Natural Sciences and Engineering Research Council (Canada) with further support from the Canada Research Chairs program. Calculations were performed on Compute Canada facilities, funded by the Canada Foundation for Innovation and partners.

## Data availability

Simulation files as well as structure, binary and trajectory files are available on request from the authors.

## Notes

### Competing Interest Statement

The authors have declared no competing interest.

## References

1. Vane J., Y. Bakhle, R. Botting. CYCLOOXYGENASES 1 AND 2. Annual review of pharmacology and toxicology. 1998;38(1):97–120.

2. Smith W. L., D. L. DeWitt, R. M. Garavito. Cyclooxygenases: structural, cellular, and molecular biology. Annual review of biochemistry. 2000;69(1):145–182.

3. Schneider C., A. Pozzi. Cyclooxygenases and lipoxygenases in cancer. Cancer and Metastasis Reviews. 2011;30(3-4):277–294.

4. Rouzer C. A., L. J. Marnett. Cyclooxygenases: structural and functional insights. J Lipid Res. 2009;50(Supplement):S29–S34.

5. Wang M.-T., K. V. Honn, D. Nie. Cyclooxygenases, prostanoids, and tumor progression. Cancer and Metastasis Reviews. 2007;26(3-4):525.

6. Furse K. E., D. A. Pratt, N. A. Porter, T. P. Lybrand. Molecular dynamics simulations of arachidonic acid complexes with COX-1 and COX-2: insights into equilibrium behavior. Biochemistry. 2006;45(10):3189–3205.

7. Furse K. E., D. A. Pratt, C. Schneider, A. R. Brash, N. A. Porter, T. P. Lybrand. Molecular dynamics simulations of arachidonic acid-derived pentadienyl radical intermediate complexes with COX-1 and COX-2: insights into oxygenation regio-and stereoselectivity. Biochemistry. 2006;45(10):3206–3218.

8. Blobaum A. L., L. J. Marnett. Structural and functional basis of cyclooxygenase inhibition. J Med Chem. 2007;50(7):1425–1441.

9. Smith W. L., Y. Urade, P.-J. Jakobsson. Enzymes of the cyclooxygenase pathways of prostanoid biosynthesis. Chem Rev. 2011;111(10):5821–5865.

10. Garavito R. M., M. G. Malkowski, D. L. DeWitt. The structures of prostaglandin endoperoxide H synthases-1 and-2. Prostaglandins & other lipid mediators. 2002;68:129–152.

11. Luong C., A. Miller, J. Barnett, J. Chow, C. Ramesha, M. F. Browner. Flexibility of the NSAID binding site in the structure of human cyclooxygenase-2. Nature structural biology. 1996;3(11):927–933.

12. Rao P., E. E. Knaus. Evolution of nonsteroidal anti-inflammatory drugs (NSAIDs): cyclooxygenase (COX) inhibition and beyond. Journal of Pharmacy & Pharmaceutical Sciences. 2008;11(2):81–110s.

13. Otto J. C., W. L. Smith. The orientation of prostaglandin endoperoxide synthases-1 and-2 in the endoplasmic reticulum. J Biol Chem. 1994;269(31):19868–19875.

14. Spencer A. G., J. W. Woods, T. Arakawa, I. I. Singer, W. L. Smith. Subcellular localization of prostaglandin endoperoxide H synthases-1 and-2 by immunoelectron microscopy. J Biol Chem. 1998;273(16):9886–9893.

15. Smith W. L., M. G. Malkowski. Interactions of fatty acids, nonsteroidal anti-inflammatory drugs, and coxibs with the catalytic and allosteric subunits of cyclooxygenases-1 and-2. J Biol Chem. 2019;294(5):1697–1705.

16. Dong L., A. J. Vecchio, N. P. Sharma, B. J. Jurban, M. G. Malkowski, W. L. Smith. Human cyclooxygenase-2 is a sequence homodimer that functions as a conformational heterodimer. J Biol Chem. 2011;286(21):19035–19046.

17. Rouzer C. A., L. J. Marnett. Endocannabinoid oxygenation by cyclooxygenases, lipoxygenases, and cytochromes P450: cross-talk between the eicosanoid and endocannabinoid signaling pathways. Chem Rev. 2011;111(10):5899–5921.

18. Corradi V., B. I. Sejdiu, H. Mesa-Galloso, H. Abdizadeh, S. Y. Noskov, S. J. Marrink, D. P. Tieleman. Emerging Diversity in Lipid–Protein Interactions. Chem Rev. 2019.

19. Marrink S. J., V. Corradi, P. C. Souza, H. I. Ingólfsson, D. P. Tieleman, M. S. Sansom. Computational modeling of realistic cell membranes. Chem Rev. 2019;119(9):6184–6226.

20. Enkavi G., M. Javanainen, W. Kulig, T. Róg, I. Vattulainen. Multiscale Simulations of Biological Membranes: The Challenge To Understand Biological Phenomena in a Living Substance. Chem Rev. 2019.

21. Blomberg L. M., M. R. Blomberg, P. E. Siegbahn, W. A. van der Donk, A.-L. Tsai. A quantum chemical study of the synthesis of prostaglandin g2 by the cyclooxygenase active site in prostaglandin endoperoxide h synthase 1. The Journal of Physical Chemistry B. 2003;107(14):3297–3308.

22. Christov C. Z., A. Lodola, T. G. Karabencheva-Christova, S. Wan, P. V. Coveney, A. J. Mulholland. Conformational Effects on the pro-S Hydrogen Abstraction Reaction in Cyclooxygenase-1: An Integrated QM/MM and MD Study. Biophys J. 2013;104(5):L5–L7.

23. Nina M., S. Bernèche, B. Roux. Anchoring of a monotopic membrane protein: the binding of prostaglandin H 2 synthase-1 to the surface of a phospholipid bilayer. European Biophysics Journal. 2000;29(6):439–454.

24. Wan S., P. V. Coveney. A comparative study of the COX-1 and COX-2 isozymes bound to lipid membranes. Journal of computational chemistry. 2009;30(7):1038–1050.

25. Fowler P. W., K. Balali-Mood, S. Deol, P. V. Coveney, M. S. Sansom. Monotopic enzymes and lipid bilayers: a comparative study. Biochemistry. 2007;46(11):3108–3115.

26. Lomize A. L., I. D. Pogozheva, M. A. Lomize, H. I. Mosberg. The role of hydrophobic interactions in positioning of peripheral proteins in membranes. BMC Struct Biol. 2007;7(1):44.

27. Balali-Mood K., P. J. Bond, M. S. Sansom. Interaction of monotopic membrane enzymes with a lipid bilayer: a coarse-grained MD simulation study. Biochemistry. 2009;48(10):2135–2145.

28. Gupta K., B. S. Selinsky, C. J. Kaub, A. K. Katz, P. J. Loll. The 2.0 Å resolution crystal structure of prostaglandin H2 synthase-1: structural insights into an unusual peroxidase. J Mol Biol. 2004;335(2):503–518.

29. Marrink S. J., H. J. Risselada, S. Yefimov, D. P. Tieleman, A. H. de Vries. The MARTINI Force Field: Coarse Grained Model for Biomolecular Simulations. The Journal of Physical Chemistry B. 2007;111(27):7812–7824.

30. Casares D., P. V. Escribá, C. A. Rosselló. Membrane lipid composition: effect on membrane and organelle structure, function and compartmentalization and therapeutic avenues. International journal of molecular sciences. 2019;20(9):2167.

31. Van Meer G., D. R. Voelker, G. W. Feigenson. Membrane lipids: where they are and how they behave. Nat Rev Mol Cell Biol. 2008;9(2): 112–124.

32. van Meer G., A. I. de Kroon. Lipid map of the mammalian cell. Journal of cell science. 2011;124(1):5–8.

33. Wassenaar T. A., H. I. Ingólfsson, R. A. Böckmann, D. P. Tieleman, S. J. Marrink. Computational lipidomics with insane: a versatile tool for generating custom membranes for molecular simulations. Journal of chemical theory and computation. 2015;11(5):2144–2155.

34. Ingólfsson H. I., M. N. Melo, F. J. Van Eerden, C. Arnarez, C. A. Lopez, T. A. Wassenaar, X. Periole, A. H. De Vries, D. P. Tieleman, S. J. Marrink. Lipid organization of the plasma membrane. J Am Chem Soc. 2014;136(41):14554–14559.

35. Corradi V., E. Mendez-Villuendas, H. I. Ingolfsson, R. X. Gu, I. Siuda, M. N. Melo, A. Moussatova, L. J. DeGagne, B. I. Sejdiu, G. Singh, T. A. Wassenaar, K. D. Magnero, S. J. Marrink, D. P. Tieleman. Lipid-Protein Interactions Are Unique Fingerprints for Membrane Proteins. Acs Central Sci. 2018;4(6):709–717.

36. Bussi G., D. Donadio, M. Parrinello. Canonical sampling through velocity rescaling. The Journal of chemical physics. 2007;126(1):014101.

37. Berendsen H. J., J. v. Postma, W. F. van Gunsteren, A. DiNola, J. R. Haak. Molecular dynamics with coupling to an external bath. The Journal of chemical physics. 1984;81(8):3684–3690.

38. Hess B., C. Kutzner, D. Van Der Spoel, E. Lindahl. GROMACS 4: algorithms for highly efficient, load-balanced, and scalable molecular simulation. Journal of chemical theory and computation. 2008;4(3):435–447.

39. Jo S., T. Kim, V. G. Iyer, W. Im. CHARMM-GUI: a web-based graphical user interface for CHARMM. Journal of computational chemistry. 2008;29(11):1859–1865.

40. Wu E. L., X. Cheng, S. Jo, H. Rui, K. C. Song, E. M. Dávila-Contreras, Y. Qi, J. Lee, V. Monje-Galvan, R. M. Venable. CHARMM-GUI membrane builder toward realistic biological membrane simulations. Journal of computational chemistry. 2014;35(27):1997–2004.

41. Jorgensen W. L., J. Chandrasekhar, J. D. Madura, R. W. Impey, M. L. Klein. Comparison of simple potential functions for simulating liquid water. The Journal of chemical physics. 1983;79(2):926–935.

42. Huang J., S. Rauscher, G. Nawrocki, T. Ran, M. Feig, B. L. de Groot, H. Grubmüller, A. D. MacKerell. CHARMM36m: an improved force field for folded and intrinsically disordered proteins. Nature methods. 2017;14(1):71–73.

43. Darden T., D. York, L. Pedersen. Particle mesh Ewald: An No log (N) method for Ewald sums in large systems. The Journal of chemical physics. 1993;98(12):10089–10092.

44. Nosé S. A molecular dynamics method for simulations in the canonical ensemble. Molecular physics. 1984;52(2):255–268.

45. Hoover W. G. Canonical dynamics: Equilibrium phase-space distributions. Physical review A. 1985;31(3):1695.

46. Parrinello M., A. Rahman. Polymorphic transitions in single crystals: A new molecular dynamics method. Journal of Applied physics. 1981;52(12):7182–7190.

47. Hess B., H. Bekker, H. J. Berendsen, J. G. Fraaije. LINCS: a linear constraint solver for molecular simulations. Journal of computational chemistry. 1997;18(12):1463–1472.

48. Schmid N., A. P. Eichenberger, A. Choutko, S. Riniker, M. Winger, A. E. Mark, W. F. van Gunsteren. Definition and testing of the GROMOS force-field versions 54A7 and 54B7. European biophysics journal. 2011;40(7):843.

49. Berendsen H. J., J. P. Postma, W. F. van Gunsteren, J. Hermans. Interaction models for water in relation to protein hydration. Intermolecular forces: Springer; 1981. p. 331–342.

50. Abraham M. J., T. Murtola, R. Schulz, S. Páll, J. C. Smith, B. Hess, E. Lindahl. GROMACS: High performance molecular simulations through multi-level parallelism from laptops to supercomputers. SoftwareX. 2015;1:19–25.

51. McGibbon R. T., K. A. Beauchamp, M. P. Harrigan, C. Klein, J. M. Swails, C. X. Hernández, C. R. Schwantes, L.-P. Wang, T. J. Lane, V. S. Pande. MDTraj: a modern open library for the analysis of molecular dynamics trajectories. Biophys J. 2015;109(8):1528–1532.

52. Sejdiu B. I., D. P. Tieleman. Lipid-Protein Interactions are a Unique Property and Defining Feature of G Protein-Coupled Receptors. Biophys J. 2020.

53. Humphrey W., A. Dalke, K. Schulten. VMD: visual molecular dynamics. Journal of molecular graphics. 1996;14(1):33–38.

54. Rose A. S., P. W. Hildebrand. NGL Viewer: a web application for molecular visualization. Nucleic Acids Res. 2015;43(W1):W576–W579.

55. Rose A. S., A. R. Bradley, Y. Valasatava, J. M. Duarte, A. Prlić, P. W. Rose. NGL viewer: webbased molecular graphics for large complexes. Bioinformatics. 2018;34(21):3755–3758.

56. Shinohara H., M. a. A. Balboa, C. A. Johnson, J. Balsinde, E. A. Dennis. Regulation of delayed prostaglandin production in activated P388D1 macrophages by group IV cytosolic and group V secretory phospholipase A2s. J Biol Chem. 1999;274(18):12263–12268.

57. Rieke C. J., A. M. Mulichak, R. M. Garavito, W. L. Smith. The role of arginine 120 of human prostaglandin endoperoxide H synthase-2 in the interaction with fatty acid substrates and inhibitors. J Biol Chem. 1999;274(24):17109–17114.

58. Vecchio A. J., B. J. Orlando, R. Nandagiri, M. G. Malkowski. Investigating Substrate Promiscuity in Cyclooxygenase-2 THE ROLE OF ARG-120 AND RESIDUES LINING THE HYDROPHOBIC GROOVE. J Biol Chem. 2012;287(29):24619–24630.

59. Tidhar R., A. H. Futerman. The complexity of sphingolipid biosynthesis in the endoplasmic reticulum. Biochimica Et Biophysica Acta (BBA)-Molecular Cell Research. 2013;1833(11):2511–2518.

60. Breslow D. K. Sphingolipid homeostasis in the endoplasmic reticulum and beyond. Cold Spring Harbor perspectives in biology. 2013;5(4):a013326.

61. Kovesi P. Good colour maps: How to design them. arXiv preprint arXiv:150903700. 2015.

62. Duncan A. L., R. A. Corey, M. S. Sansom. Defining how multiple lipid species interact with inward rectifier potassium (Kir2) channels. Proceedings of the National Academy of Sciences. 2020.

63. Fagone P., S. Jackowski. Membrane phospholipid synthesis and endoplasmic reticulum function. J Lipid Res. 2009;50(Supplement):S311–S316.

64. Marrink S. J., A. H. De Vries, A. E. Mark. Coarse grained model for semiquantitative lipid simulations. The Journal of Physical Chemistry B. 2004;108(2):750–760.

65. Dong L., H. Zou, C. Yuan, Y. H. Hong, D. V. Kuklev, W. L. Smith. Different fatty acids compete with arachidonic acid for binding to the allosteric or catalytic subunits of cyclooxygenases to regulate prostanoid synthesis. J Biol Chem. 2016;291(8):4069–4078.

66. Barreto-Ojeda E., V. Corradi, R.-X. Gu, D. P. Tieleman. Coarse-grained molecular dynamics simulations reveal lipid access pathways in P-glycoprotein. Journal of General Physiology. 2018;150(3):417–429.

67. Orlando B. J., D. R. McDougle, M. J. Lucido, E. T. Eng, L. A. Graham, C. Schneider, D. L. Stokes, A. Das, M. G. Malkowski. Cyclooxygenase-2 catalysis and inhibition in lipid bilayer nanodiscs. Archives of biochemistry and biophysics. 2014;546:33–40.

68. Yuan C., R. S. Sidhu, D. V. Kuklev, Y. Kado, M. Wada, I. Song, W. L. Smith. Cyclooxygenase allosterism, fatty acid-mediated cross-talk between monomers of cyclooxygenase homodimers. J Biol Chem. 2009;284(15):10046–10055.

69. Dong L., C. Yuan, B. J. Orlando, M. G. Malkowski, W. L. Smith. Fatty acid binding to the allosteric subunit of cyclooxygenase-2 relieves a tonic inhibition of the catalytic subunit. J Biol Chem. 2016;291(49):25641–25655.

