## Supplementary figures for "COX-1 – lipid interactions: arachidonic acid, cholesterol, and phospholipid binding to the membrane binding domain of COX-1"

### SUPPLEMENTARY INFORMATION

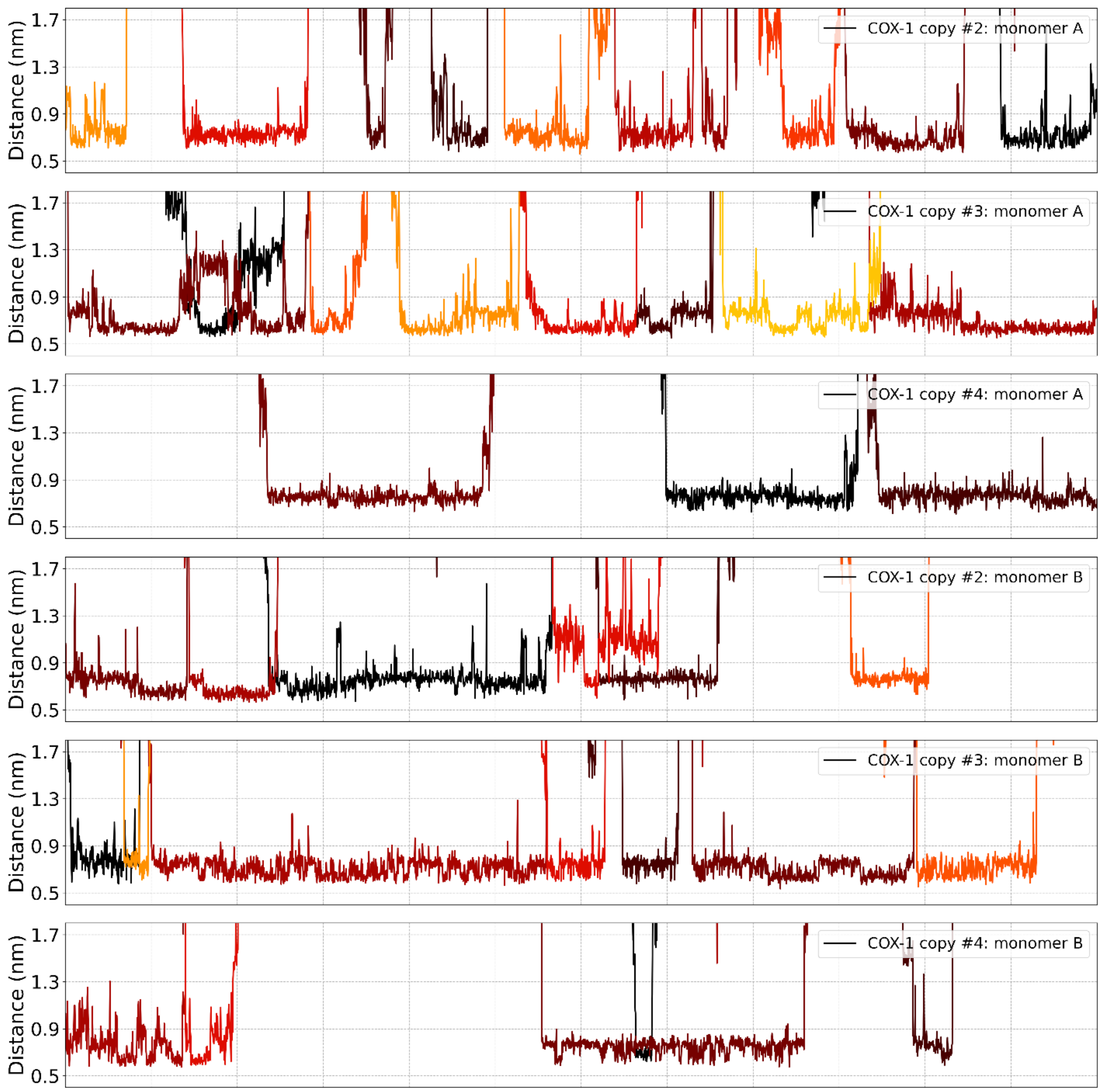


***Figure S1. Cholesterol binding to the MBD of COX-1 in CG simulations.*** *Data are for the MBDs of the rest of the proteins in the system (proteins #2, #3, and #4), to complement Figure 4 in the main text (which shows data for protein #1).*

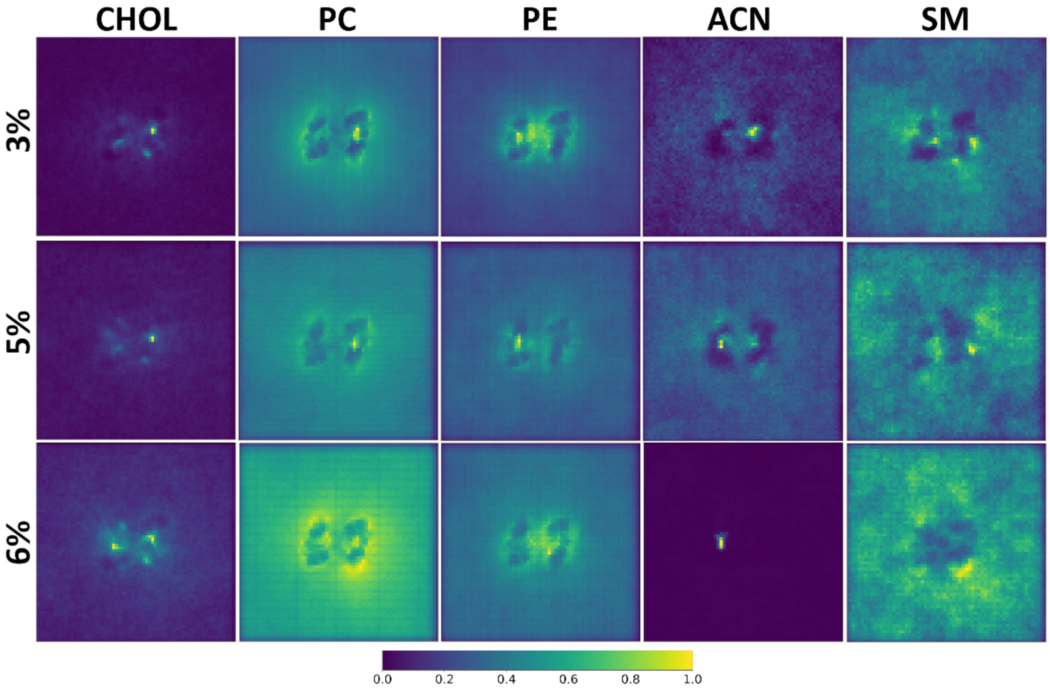


***Figure S2. 2D density profiles for ER-like membrane simulations.*** *For each lipid component the density profile for the upper leaflet is shown. In most cases, we see that the highest localization of lipids corresponds to the cavity of the MBDs. Normalization is done on a per lipid basis (and not across all lipids). Notice the creation of the 'coronal layer' around COX-1 for PC and PE lipids, closely resembling the FS layers from Figure 2.*

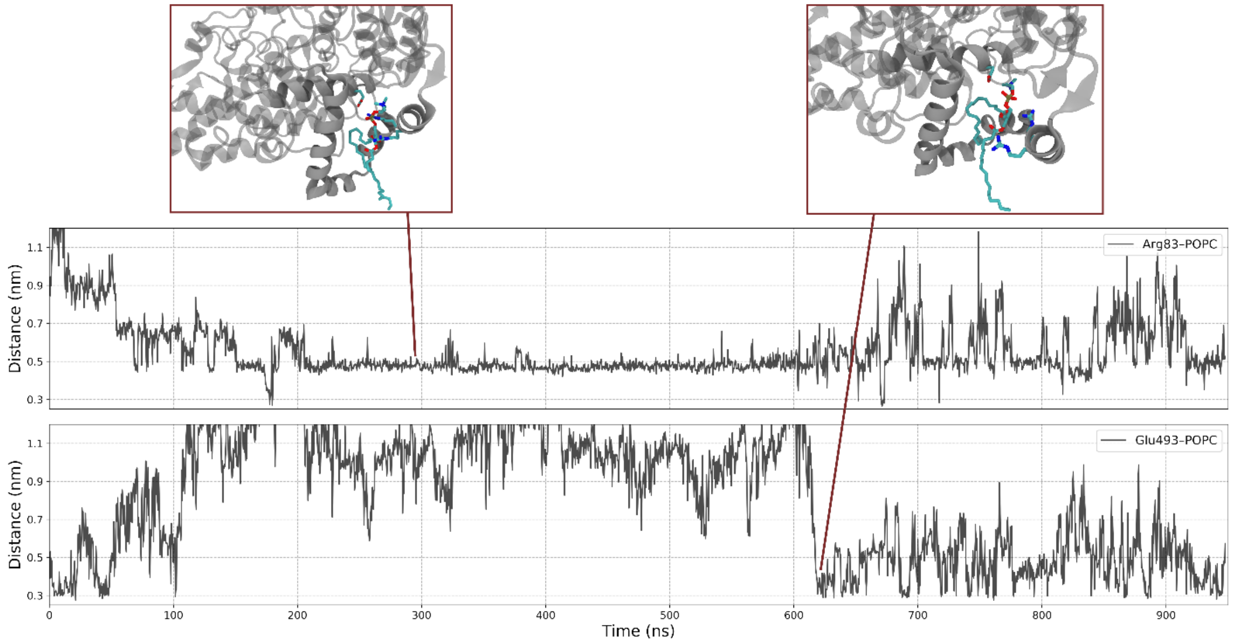


***Figure S3. POPC lipid binding to the MBD of COX-1 in AA simulations.*** *Binding of POPC to the MBD of COX-1 utilizes the same cavity within the MBD, however the binding itself differs. First, POPC binding is more erratic owing to its higher flexibility within the site. The distances shown here are between the P atom of bound POPC lipid and either the sidechain carboxyl C atom (for Glu-493 distance) or guanidino carbon atom (for Arg-83 distance)*

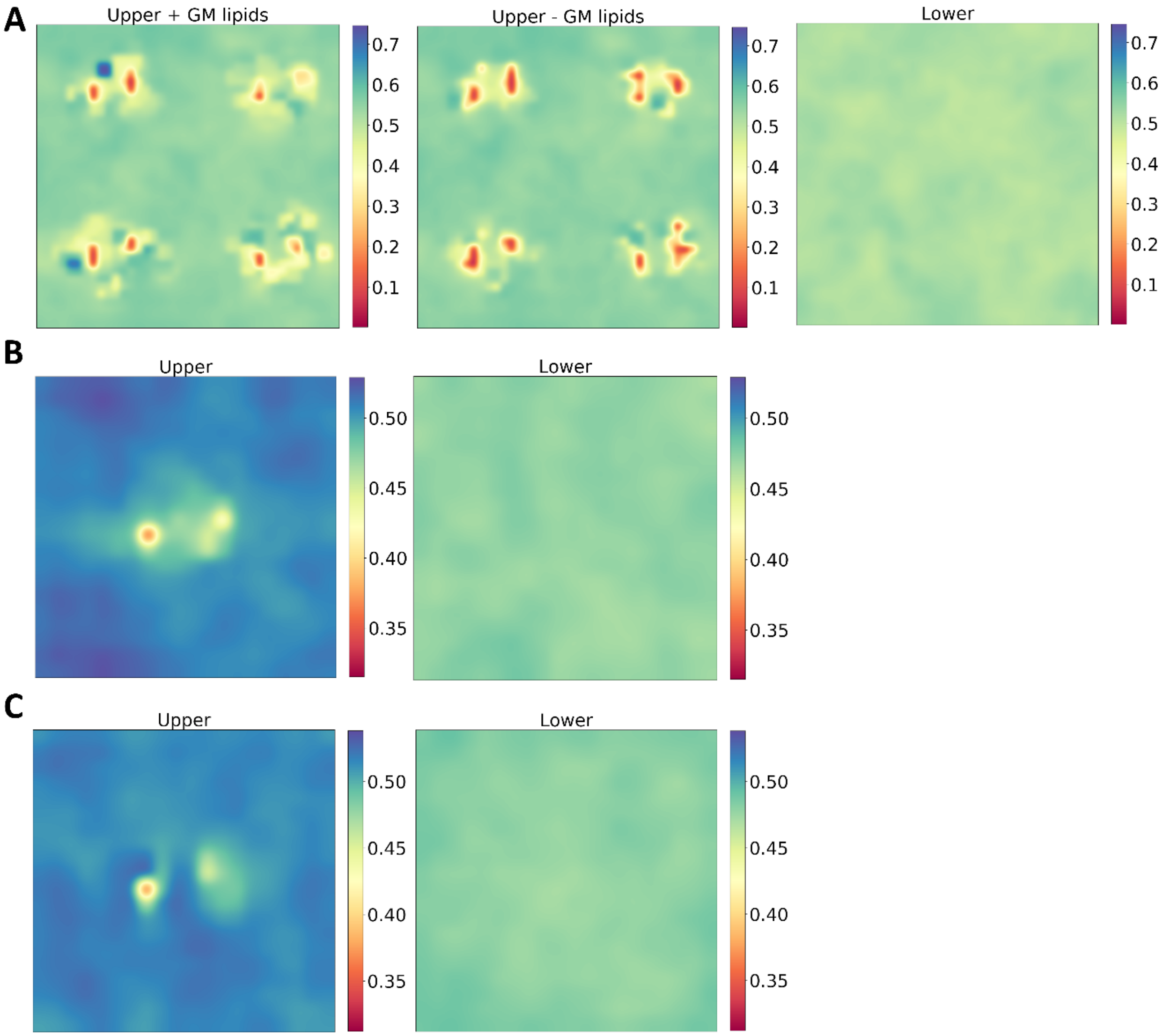


***Figure S4. Coarse-Grained order parameters for some multicomponent systems.*** *Data are for the complex membrane setup (**A**), 6% cholesterol (**B**) and 3% cholesterol (**C**) content systems. We see a noticeable disorder caused by the protein caused on the upper leaflet in all systems, yet we do not see any change on the lower leaflet of the membrane.*

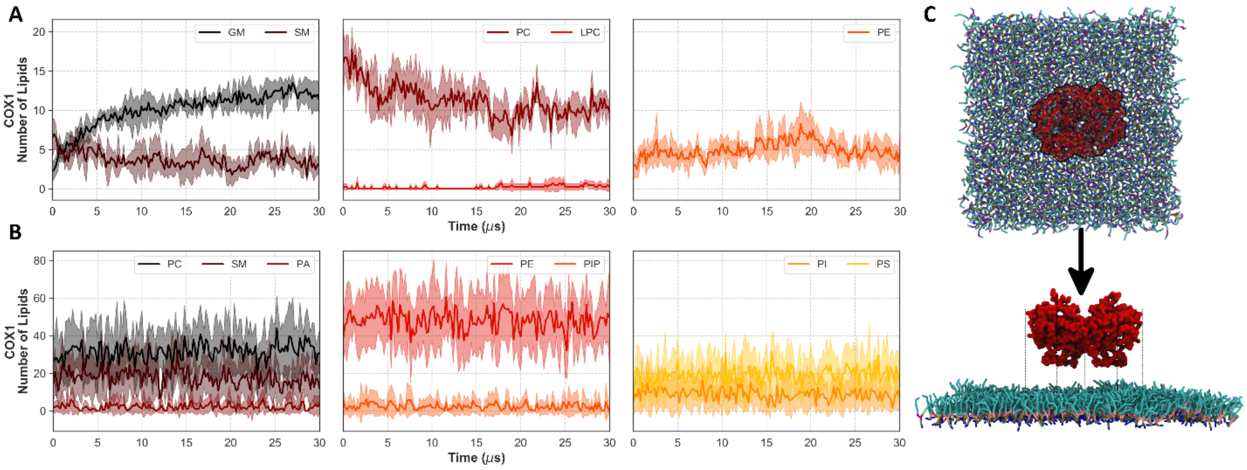


***Figure S5. Lipid count in the close proximity of embedded COX-1 proteins in the complex system.*** *Lipids are grouped according to their headgroup type into the following categories: PC, PE, PS, GM, SM, PA, PI, PIP, LPC.* ***A.*** *Lipid count within a 7Å radius around proteins.* ***B.*** *Lipid count within the perimeter formed by the projection of the outermost residues of the protein.* ***C.*** *Description of how lipids are counted for the lower leaflet. The largest circumference drawn along the outermost points of the protein (colored in black) is projected onto the lower leaflet to define the area for counting lipids. Lipids that fall within that surface are counted and displayed in* ***B****.*

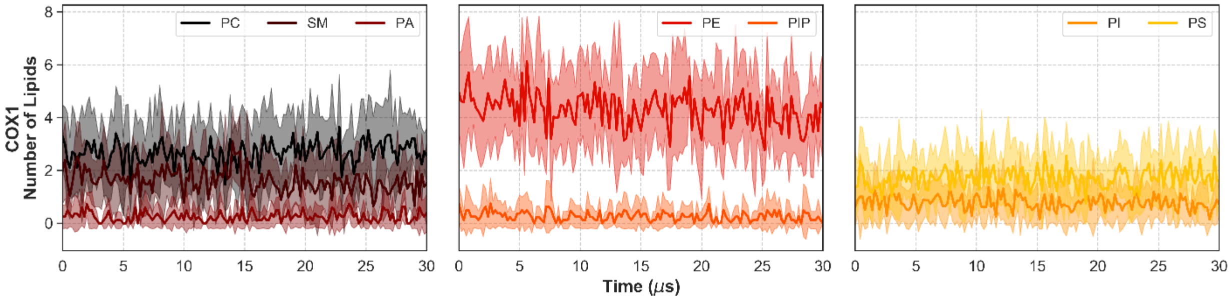


***Figure S6. Lipid count within 7Å of COX-1 in the complex membrane system.*** *Data are shown for the lower leaflet only. Calculations are done similar to Figure S5, but here the MBDs of the proteins are projected into the lower leaflet to define the surface for counting lipids. Only lipid species that fall within that surface are counted.*

***Table S1. Detailed overview of simulated systems. For each system we show the lipid composition as well as the total simulation time.***

| Resolution | Description | Composition (%) | Simulation time (μs) |
|---|---|---|---|
| CG | Complex Membrane Setup / Plasma membrane | See **Table S2** *(.xlsx)* | 30 |
| CG | ER-like membrane setup w/ 3% cholesterol | See **Table S3** *(.xlsx)* | 10 |
| | | See **Table S3** *(.xlsx)* | |
| CG | ER-like membrane setup w/ 5% cholesterol | See **Table S3** *(.xlsx)* | 4 |
| | | See **Table S3** *(.xlsx)* | |
| CG | ER-like membrane setup w/ 6% cholesterol | See **Table S3** *(.xlsx)* | 10 |
| | | See **Table S3** *(.xlsx)* | |
| CG | Large POPC only | POPC (1141 lipids) | 10 |
| AA | Low CHOL | CHOL:13;POPC:87 | 1 |
| AA | High CHOL | CHOL:40; POPC:60 | 0.45 |
| AA | 40% Arachidonic Acid (ARAN) | ARAN:40;POPC60 | 0.95 |
| AA | 40% Arachidonic Acid (ARANP) | ARANP:40;POPC60 | 0.95 |
| AA | Large POPC membrane | POPC:100 (1140 lipids) | 0.2 |
| AA | Large POPC membrane / surface tension: 50 | POPC:100 (1140 lipids) | 0.275 |
| AA | Large POPC membrane / surface tension: 300 | POPC:100 (1140 lipids) | 0.25 |
| AA | Large POPC membrane / GROMOS54A7 ff | POPC:100 (1140 lipids) | 0.2 |
| AA | Large POPC membrane / histidine protonation state | POPC:100 (1140 lipids) | 0.075 |
| AA | Reference POPC bilayer / no protein | POPC:100 (1140 lipids) | 0.2 |
